# Fluctuating selection breaks Hamilton’s rule for the evolution of altruism

**DOI:** 10.1101/2025.04.18.649474

**Authors:** Feng Zhang, Alan Hastings, Cang Hui

## Abstract

Altruistic behaviours, those that benefit recipients at a cost to donors, have long posed a puzzle in evolutionary biology and sociobiology. Established theories, such as kin selection, group selection, and reciprocal altruism, explain altruism via assortment mechanisms that depend on preferential interactions among altruists to ensure a higher average payoff than selfish individuals. Hamilton’s rule defines the minimum level of such assortative interactions required for altruism to evolve. However, in populations with limited growth, increased competition can erode the selective advantage of altruism and thus require more stringent Hamilton’s rule. Here, we propose a fundamentally different mechanism: fluctuating selection driven by increased competition due to added social benefits of altruism in populations with limited carrying capacity. Under fluctuating selection, altruists can invade a selfish population without the need of assortment mechanisms and outcompete selfish individuals through a transient phase, even when the cost of altruism exceeds its direct benefit. Classical invasion analysis, which compares the long-term growth rates of rare altruistic mutants and resident selfish individuals, fails in our model because both strategies exhibit zero long-term growth. Instead, we show that altruism can invade when its short-term, time-averaging growth rate exceeds that of selfish individuals, which necessitates a prolonged transient phase. This invasion is enabled by payoff-regulated density dependence, which captures the dynamic interplay between payoff and density on fitness under fluctuating selection. Our findings challenge the necessity of Hamilton’s rule and show that genuine altruism, with less average payoff than selfish individuals, even extremely self-sacrificial, can emerge under natural selection. This suggests that altruism may arise not from preferential interaction, but as an adaptive response to population fluctuations in constrained environments, providing an alternative paradigm for understanding the evolution of altruistic behaviour.

## Introduction

The evolution of altruistic behaviour in a selfish population has been a longstanding question in evolutionary biology and sociobiology [1-5]. It is now accepted that assortment—preferential interactions between altruists—is essential for the evolution of altruism [6-9]. With solely random interactions, altruists cannot invade and persist in a selfish population, as they would consistently achieve lower average payoffs than selfish individuals. As a result, current theories—including kin selection [10-12], group selection [13, 14], and reciprocal altruism [15-17]—focus on mechanisms that establish assortative interactions to boost the average payoff of altruists over that of selfish individuals, typically through genetic, spatial or temporal affinity among altruists. Hamilton’s rule, in its various forms, identifies the minimum intensity of such assortments for altruism to evolve, ensuring that altruists receive higher average payoffs than selfish individuals [18], a concept that can be described as ‘narcissistic altruism’ from the perspective of selfish gene theory [19].

Although assortment mechanisms promote altruism, added benefits of altruism can also intensify competition among individuals, potentially reducing or even negating the selective advantage of altruism, especially in constrained environments with limited carrying capacities [20-24]. In contrast, we here demonstrate that heightened competition resulting from added social benefits of altruism can trigger fluctuating selection—where the strength and direction of selection swing with the wax and wane of population fluctuations [25-28]. This enables altruists to invade and dominate a selfish population without relying on assortment mechanisms or adherence to Hamilton’s rule, although altruists consistently receive a lower average payoff than selfish individuals. Moreover, altruism can emerge from selfishness even when the cost of altruism exceeds its resulting benefit, leading to the behaviour of extreme self-sacrifice. This shows that, instead of hindering the evolution of altruism, the intensified competition from social benefits plays a crucial role in fostering altruism to emerge and establish particularly in fluctuating environments.

### Regime shift for altruism evolution

Consider a population where individuals interact altruistically or selfishly. An altruist incurs a cost *c* to provide a benefit *b* to its opponent, while a selfish individual pays no cost and provides no benefit. An altruist thus receives a payoff of *b*−*c* when interacting with another altruist, and −*c* when interacting with a selfish individual. A selfish individual receives a payoff of *b* when interacting with an altruist and zero when interacting with another selfish individual. When *b* > *c* > 0, this describes a Prisoner’s Dilemma (PD) game [29, 30], where selfishness is the Nash equilibrium, even though mutual altruism yields the highest overall payoff (Pareto optimal), thus creating a social dilemma. When *c* > *b* > 0, altruism represents extreme self-sacrifice as mutual altruism yields the lowest overall payoff, and selfishness remains the Nash equilibrium. If individuals interact with each other at random (without assortment), the average payoffs for altruists and selfish individuals are *ξ* = *pb* − *c* and *ζ* = *pb*, respectively, where *p* is the frequency of altruists in the population. Notably, an altruist’s average payoff is always lower than that of a selfish individual (*ξ* < *ζ*).

We assume that individuals are identical except for their behavioural strategies; the growth rate of each strategy depends additively on the average payoff obtained by individuals of that strategy; and the population’s carrying capacity is determined by external factors and independent of the payoffs of the strategies. Thus, the dynamics of both strategies can be described by the Ricker model [31, 32]:

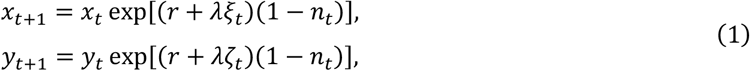

where *x*_*t*_ and *y*_*t*_ represent the densities of altruists and selfish individuals at generation *t*, respectively; *r* is the intrinsic growth rate of the population; *λ* (> 0) is the selection intensity on payoffs; *n*_*t*_ = *x*_*t*_ *+ y*_*t*_ is the total population density at generation *t*, and (1 − *n*_*t*_) expresses the density-dependent regulation on population growth (with the carrying capacity rescaled to one). The terms in square brackets are the relative growth rates of altruistic and selfish individuals at generation *t*, i.e., log(*x*_*t+*1_/*x*_*t*_) and log(*y*_*t+*1_/*y*_*t*_). The relative growth rates can be decomposed into two parts: terms (*r + λξ*_*t*_) and (*r + λζ*_*t*_) represent the sum of the intrinsic growth rate and the additive effect of payoff, as in the classical models of evolutionary game theory [33-37], while terms −(*r+ λξ*_*t*_)*n*_*t*_ and −(*r+ λζ*_*t*_)*n*_*t*_ are the payoff-regulated density dependence and describe intensified competition from payoff increase.

The discrete replicator equation [38] can be recovered from Eq. (1) if we remove the density-dependent term (1 − *n*_*t*_) (see Supplementary Material S1). This model, therefore, extends the classic evolutionary game theory by incorporating density-dependent regulation on population growth. Density dependence can introduce complex population dynamics [39, 40], leading to fluctuating selection [25-27], where altruists may experience either a lower or higher relative growth rate than selfish individuals, depending on whether the population density is increasing (1 − *n*_*t*_ > 0) or decreasing (1 − *n*_*t*_ < 0). Since altruists have a lower average payoff than selfish individuals (*ξ*_*t*_ < *ζ*_*t*_), the density of altruists increases slower than that of selfish individuals during periods of population rise [0 < log(*x*_*t+*1_/*x*_*t*_) < log(*y*_*t+*1_/*y*_*t*_)] while it also decreases less rapidly during population decline [0 > log(*x*_*t+*1_/*x*_*t*_) > log(*y*_*t+*1_/*y*_*t*_)]. This model thus captures the evolutionary dynamics of altruism under fluctuating selection [25-27], driven by the payoff-regulated density dependence and the intrinsic instability of the system (see models with other stochastic factors in Supplementary Materials S3 and S4).

In a population consisting entirely of selfish individuals (*x* = 0), the Ricker model simplifies to *y*_*t+*1_ = *y*_*t*_ exp[*r*(1 − *y*_*t*_)], where *ζ*_*t*_ = 0 and *n*_*t*_ = *y*_*t*_. The dynamics in this case are governed solely by the parameter *r* [39, 40]: if 0 < *r* ≤ 2, the population density will asymptotically approach a stable equilibrium (*n*^∗^ = 1); however, if *r* > 2, the equilibrium becomes destabilized, causing the population to fluctuate periodically or chaotically around a long-term average density that is equal to the equilibrium (*ñ* = *n*^∗^). For altruism to invade a selfish population, these fluctuations are necessary. Rare altruistic mutants cannot invade a stable population (0 < *r* 2), but when the population fluctuates (*r* > 2), altruism can invade and eventually dominate the population, leading to the extirpation of selfish individuals after a prolonged transient phase (see Fig. 1 and 2, Fig. S2-S4). Notably, this occurs while the average payoff of the altruists stays persistently lower than that of selfish individuals (*ξ*_*t*_ < *ζ*_*t*_ for all *t*; Fig. 1C, Fig. S2C, Fig. S3C), and the same is true even if the cost of altruism exceeds the benefit (for extremely self-sacrificial altruistic behaviour; Fig. S3 and S4). Below, the invasion criterion of altruism under fluctuating selection and the transient phase during which the system regime shifts from selfishness to altruism are explained.

**Fig. 1.**
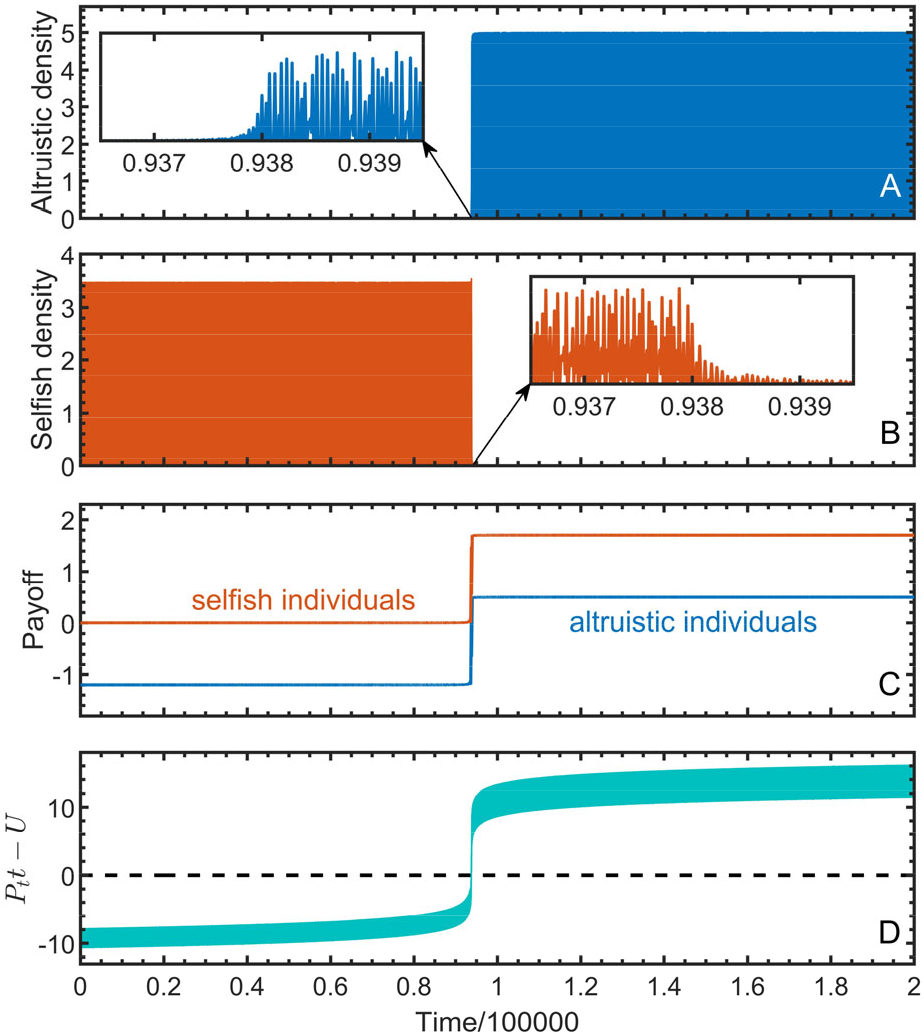
Invasion of altruism through a regime shift after a long transient phase in a chaotically fluctuating population. Panel (A) displays the dynamics of altruists starting from a small initial density in a selfish population. Panel (B) illustrates the dynamics of selfish individuals. Panel (C) shows the average payoffs of altruistic and selfish individuals. Panel (D) demonstrates how the duration of the transient phase results from the inequality (3), where *P*_*t*_*t* and *U* are the left- and right-hand side of the inequality, respectively. The parameters used are: *b* = 1.7, *c* = 1.2, *λ* = 1, *r* = 3.5, *x*_0_ = 0.00001, and *y*_0_ = 1.

**Fig. 2.**
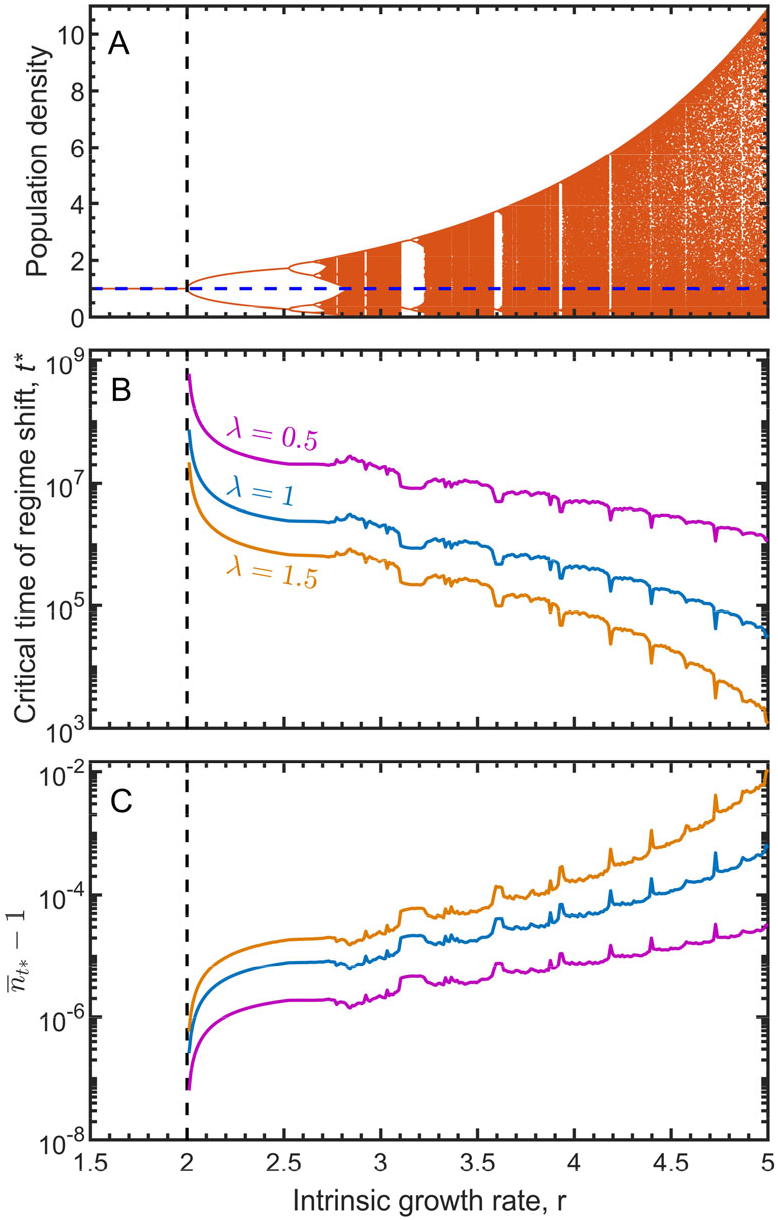
The regime shift for altruism invasion occurs in the presence of periodic and chaotic population dynamics, depending on the intrinsic growth rate of the population. Panel (A) shows the bifurcation diagram of selfish population dynamics (*y*_*t+*1_ = *y*_*t*_ exp[*r*(1 − *y*_*t*_)]), with the horizontal dashed line indicating the long-term average population density. Panel (B) shows the critical time of regime shift. Panel (C) shows 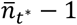, corresponding to the curves on panel (B), where 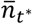 is the average population density over generations up to the critical time *t*^∗^ (transient phase of regime shift). The parameters used are: *b* = 1, *c* = 0.6, *x*_0_ = 0.00001, and *y*_0_ = 1.

### Invasion criterion and transient phase

The invasion of a rare mutant strategy in variable environments requires its long-term growth rate (i.e., the Lyapunov exponent, defined as the limit of the average relative growth rate over *t* generations as *t* approaches infinity) to be greater than that of the resident population [41, 42]. However, in our model, the long-term growth rates of both initially rare altruists and the pre-invasion selfish population are zero (see Supplementary Material S2). In such cases, if the initial altruists increase relative to selfish individuals in terms of the short-term average growth rate over *t* generations, altruism can invade selfishness in the population.

From Eq. (1), the density ratio of altruists to selfish individuals can be derived as *x*_*t+*1_/*y*_*t+*1_ = (*x*_*t*_/*y*_*t*_)exp [*λ*(*ξ*_*t*_ − *ζ*_*t*_)(1 − *n*_*t*_)]. Note that the term in square brackets is the change rate of the density ratio, which represents the difference in relative growth rate between altruists and selfish individuals (see Supplementary Material S2). Consequently, altruists increase relative to selfish individuals when the average change rate of the density ratio over *t* generations is positive. This leads to the invasion condition of altruism in a selfish population (see Supplementary Material S2 for more details):

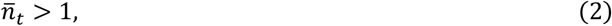

where 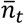 is the average population density over the *t* generations. This invasion criterion indicates that for altruists to successfully invade a selfish population, the total population density must, on average, fluctuate above one before the regime shift occurs (Fig. 3). This threshold corresponds to the long-term average population density prior to the introduction of altruism (Fig. 2A; Supplementary Material S2).

**Fig. 3.**
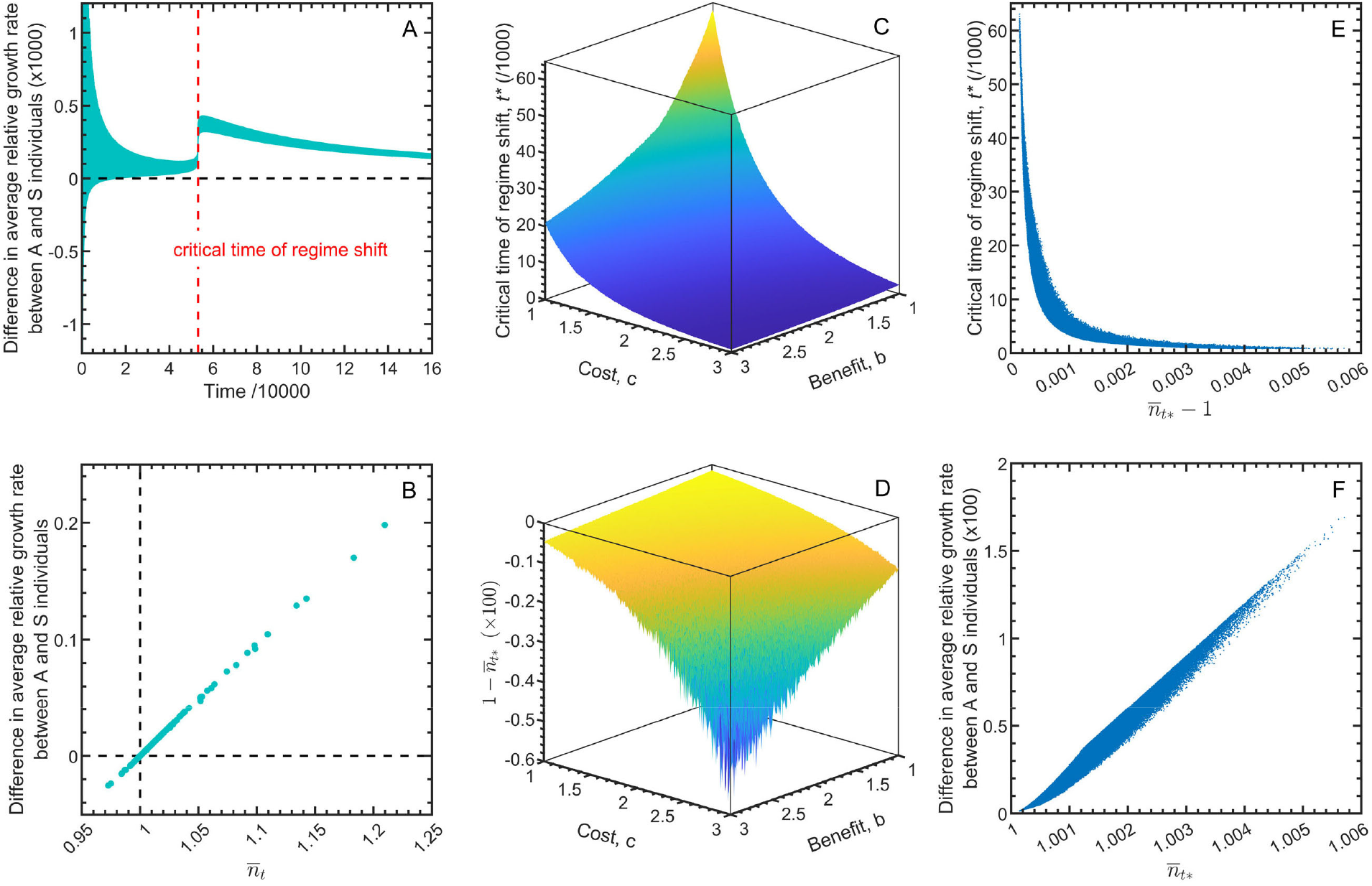
Invasion criterion for altruism in fluctuating selfish populations, with 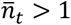. Panel (A) shows the variation of the difference in average relative growth rate between altruistic (A) and selfish (S) individuals over past *t* generations during the process of altruism invasion. Panel (B) shows that these growth differences depend on the average population density over past *t* generations, 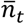. Panels (C) and (D) show the critical time of the regime shift for altruism (i.e., transient phase) and the level of 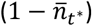 in relation to the cost and the benefit, where 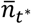 sis the average population density during the transient phase. Panel (E) shows that the critical time is inversely proportional to 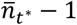. Panel (F) shows that the differences in average relative growth rate between altruistic and selfish individuals during the transient phase depend on 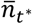. For (A) and (B), the parameters used are *b* = 1.5, *c* = 1, *λ* = 1, *r* = 4, *x*_0_ = 0.00001, and *y*_0_ = 1. For (C) to (F), both *b* and *c* are assigned equally spaced values from 1 to 3, resulting in a total of 300 × 300 combinations (whether *b* > *c* or *b* < *c*), *λ* = 1, *r* = 3, *x*_0_ = 0.0001, and *y*_0_ = 1.

Two factors are responsible for the fulfillment of the invasion criterion. First, in the Ricker model (Eq. 1), the long-term average population density of a pre-invasion selfish population always converges to one (*ñ* = *n*^∗^ = 1), regardless of whether the population fluctuates (Fig. 2A; Supplementary Material S2). However, the average population density over past *t* generations can still exceed the long-term attractor (i.e., 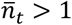) intermittently (see cyan-colored time ranges in Fig. 4A-C). These intermittent periods occur when the population density is below the initial density (i.e., *n*_*t*_ < *n*_0_) because 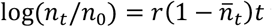 for the pre-invasion population composed purely of selfish individuals. Second, with the initial introduction of altruism, the initial density of altruists further dwindles while the density of selfish individuals increases (Fig. 5A-B), with the average population density overcompensated to rise above one (Fig. 4D-I, Fig. 5C). This fluctuation in population density, averaging above one over *t* generations, satisfies the invasion criterion (inequality 2), ensuring the successful invasion of altruism (see Fig. 2B and C, Fig. 3-5, Fig. S4B and C).

**Fig. 4.**
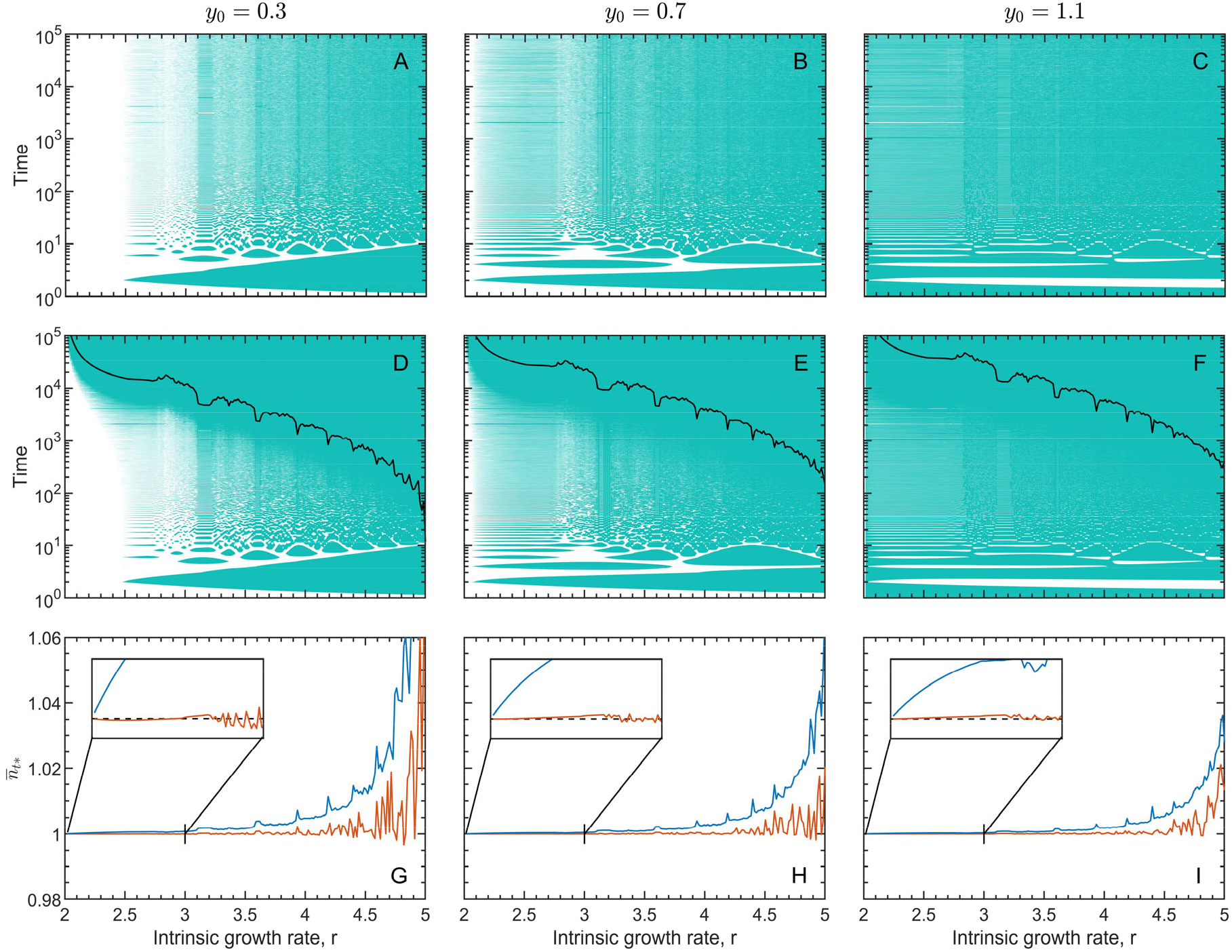
The average population density over past *t* generations under different intrinsic growth rates. Top and middle panels show the average population density changing over time (along the vertical axis) without (A-C) versus with (D-F) altruism invasion. Cyan color indicates that the average population density at the time is greater than one, white color indicates below than one. The blank lines on panels (D-F) indicate the critical time of regime shift. Panels (G)-(I) show the average population density during the transient phase, 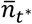, without (red) or with (blue) altruism invasion. The parameters used for (D)-(I) are *b* = 1, *c* = 0.7, *λ* = 1 and *x*_0_ = 0.0005.

**Fig. 5.**
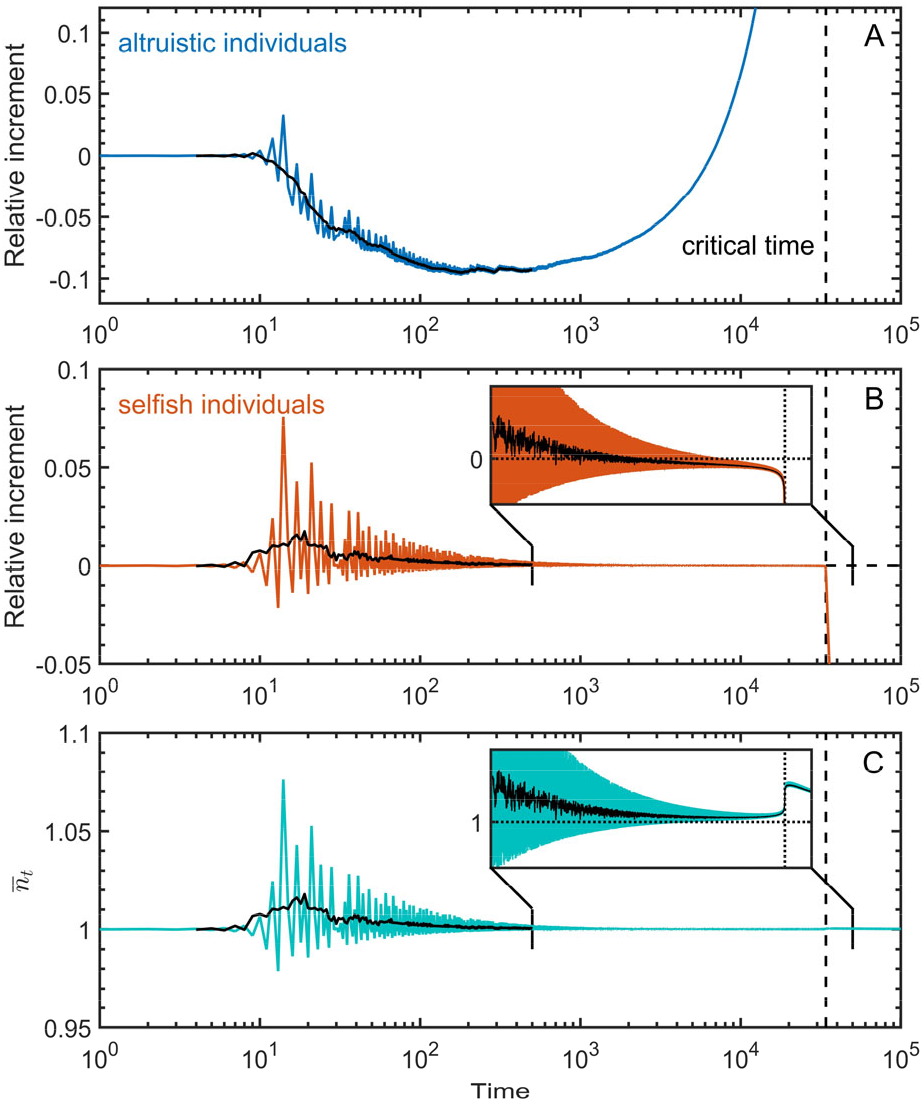
The inflation of average population density due to selfish individuals exploiting initial altruists. Relative increment refers to the difference between time-averaging and initial densities divided by initial density. Panel (A) and (B) show how the relative increments of altruistic and selfish individuals change over time. Panel (C) illustrates the dynamics of time-averaging population density, 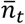, over time. Colored lines represent these quantities, while black lines are their moving averages forward and backward three timesteps. The parameters used are: *b* = 1.5, *c* = 1.09, *λ* = 1, *r* = 3, *x*_0_ = 0.0001 and *y*_0_ = 1.

To determine the duration of the transient phase (*t*^∗^) for altruism invasion, we phenomenologically examine the density ratio *x*_*t*_/*y*_*t*_. Let *x*_*t*_/*y*_*t*_ > 1 signal that altruists have taken over the population after the transient phase, which is equivalent to the following inequality (see Supplementary Material S2):

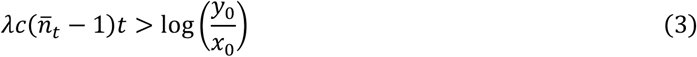

where *x*_0_ and *y*_0_ are the initial densities of altruists and selfish individuals. Given the initial rarity of altruists, we have *y*_0_/*x*_0_ > 1 and thus a positive logarithm of the initial density ratio between selfish and altruistic individuals *U* ≡ log(*y*_0_/*x*_0_) > 0. Therefore, for inequality (3) to hold, it is necessary that 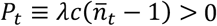, which is consistent with the invasion criterion of altruism (inequality 2). Notably, *P*_*t*_ represents the difference in the time-averaging relative growth rate between altruists and selfish individuals over the *t* generations (see Supplementary Material S2). Inequality (3) thus implies that altruists will take over when the difference in relative growth rate between altruists and selfish individuals accumulated over *t* generations (*P*_*t*_*t*) exceeds the logarithm of the initial density ratio (*U*), i.e. *P*_*t*_*t* > *U*. The boundary case, 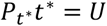, provides an estimate for the duration of the transient phase (*t*^∗^), supported by numerical simulations (Fig. 1D, panel D of Fig. S2 and S3).

Letting 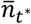 be the average population density during the transient phase, the transient phase 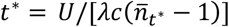 is feasible only if 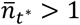 (Fig. 2B and C, Fig. 3C-F, Fig. S4B and C), and is inversely proportional to 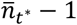 (Fig. 3E). As the population mainly consists of selfish individuals during the transient phase, the average population density is predominantly influenced by these selfish individuals (Fig. 5). The higher the average population density (i.e., the larger 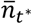), the sooner the selfish individuals are extirpated (Fig. 2B and C, Fig. 3, Fig. S4B and C): selfishness effectively becomes its own undoing.

Population fluctuations driven by environmental stochasticity (see Supplementary Material S3) and demographic stochasticity (see Supplementary Material S4), rather than intrinsic instability, can also trigger the regime shift from selfishness to altruism (Fig. S5 and S7, Fig. S9C and S9D). While population fluctuations are a necessary condition, they are not sufficient; the average population density must also exceed the threshold of invasion criterion (inequality 2; Fig. 3, Fig. S5C and S5F). If the average density of a fluctuating population remains below the threshold, altruism invasion cannot be guaranteed (Fig. S6, Fig. S9E and S9F). Additional simulations confirm that altruism invasion also occurs in the discrete logistic model, where growth factors (*x*_*t+*1_/*x*_*t*_ and *y*_*t+*1_/*y*_*t*_) are linear (see Supplementary Material S5, Fig. S10), suggesting that the Ricker model is not essential for altruism invasion. Below, we extend the invasion of altruism for any model forms under fluctuating selection.

### A generalized theory

From Eq. (1), the relative growth rates of altruists and selfish individuals can be rewritten as log(*x*_*t+*1_/*x*_*t*_) = (*r + λξ*_*t*_) − (*r+ λξ*_*t*_)*n*_*t*_ and log(*y*_*t+*1_/*y*_*t*_) = (*r + λζ*_*t*_) − (*r+ λζ*_*t*_)*n*_*t*_. Evidently, the first terms on the right-hand side represent density-independent growth rate additively depending on the average payoffs and the second terms describe payoff-regulated density dependence. Accordingly, a generalized model for the evolution of altruism can be formulated as

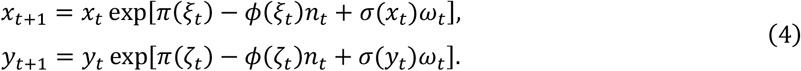

Here, *π*(·) and *ϕ*(·) are positive and increasing functions of payoffs; the former represents the density-independent growth rate, which is additively influenced by the average payoff, while the later describes payoff-regulated competition intensity, which underlies density dependence. The terms *σ*(*x*_*t*_)*ω*_*t*_ and *σ*(*y*_*t*_)*ω*_*t*_ represent environmental stochasticity when the noise intensities *σ*(*x*_*t*_) and *σ*(*y*_*t*_) are constant, and demographic stochasticity when 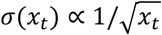 and 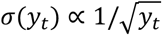, with *ω*_*t*_ following a standard normal distribution. In particular, when *ϕ*(·) is constant, density dependence is not affected by payoff, and Eq. (4) reduces to a model in which payoffs only influence population growth additively, as in the classical models of evolutionary game theory [33-37]. However, as shown below, when *ϕ*(·) increases with payoff, this payoff-regulated density dependence critically determines the evolution of altruism.

As altruists are rare before the regime shift from selfishness to altruism, the expected difference in relative growth rate between altruists and selfish individuals, i.e., 𝔼[log(*x*_*t+*1_/*x*_*t*_) − log(*y*_*t+*1_/*y*_*t*_)], can be approxima*t*ed as [*π*(−*c*) − *π*(0)] − [*ϕ*(−*c*) − *ϕ*(0)]*n*_*t*_, the sign of which depends on population density *n*_*t*_. Under fluctuating selection, altruists can invade a selfish population if the expectation of their time-averaging relative growth rate over *t* generations is greater than that of selfish individuals, i.e., 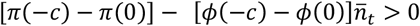, which leads to the following invasion condition

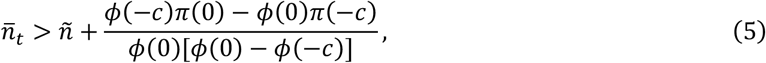

where *ñ* = *π*(0)/*ϕ*(0) is the long-term average density of the pre-invasion selfish population. Evidently, when the last term (the fraction) in inequality (5) is negative, altruists can more easily invade the selfish population. In contrast, when function *ϕ*(·) is constant, the right-hand side of inequality (5) becomes infinity, making the inequality impossible to hold and thus the un-invadability of the selfish population by altruism. This indicates that increased competition intensity from rising payoffs, captured by the increasing function *ϕ*, provides an alternative mechanism for the evolution of altruism.

As an example, in model (4), let *π*(*ξ*) = *α+ βξ* and *ϕ*(*ξ*) = *γ+ μξ*, with *α, β, γ* and *μ* being positive parameters. The difference in relative growth rate between altruists and selfish individuals becomes −*c*(*β* − *μn*_*t*_). This implies that altruists can have a higher or lower growth rate than selfish individuals depending on whether *n*_*t*_ > *β*/*μ*, leading to fluctuating selection. According to inequality (5), the condition for the evolution of altruism is 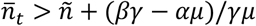 with *ñ* = *α*/*γ*. If ∂*ϕ*/∂*ξ* = *μ* > *βγ*/*α* (i.e. the sensitivity of function *ϕ* to the change of *ξ* is greater than a positive threshold), altruism can more easily invade the selfish population. In the Ricker model (Eq. 1), *ñ* = 1 and the last term in inequality (5) is zero, and thus inequality (5) becomes 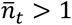, confirming the invasion condition of inequality (2). Overall, an increasing function *ϕ* of payoffs, representing increased competition of density-dependent regulation induced by added benefits of altruism, can result in fluctuating selection in systems with limited carrying capacity. Such a payoff-regulated density dependence allows altruists to invade a selfish population without the need for assortment mechanisms, thus free from Hamilton’s rule.

## Discussion

Fluctuating selection, referring to changes in the strength and direction of selection over time, is commonly driven by environmental changes, such as seasonal variations, resource availability, or interactions with other species [25-27]. This concept is frequently used to explain the maintenance of genetic variation in variable environments [25-28]. Here, we show that the payoff-regulated density dependence of population growth (i.e., increased competition from added payoffs in systems with limited carrying capacity) can induce fluctuating selection, allowing altruism to evolve in selfish populations even in the absence of assortment mechanisms (Fig. 1 and 2, Fig. S2-S4). Population fluctuations can be triggered by system instability, environmental stochasticity, or demographic stochasticity. These results underscore the role of interaction between payoff and population density in driving fluctuating selection, distinct from the effects of variable environments on selection process typically considered in population genetics [25-28]. Enhanced density-dependent regulation, driven by intensified competition from the social benefits of altruism, can trigger fluctuating selection, a key force in the evolution of altruism. This underscores the cross effect of payoffs and population density (*ξ*_*t*_*n*_*t*_) on population growth. In contrast, classical models of evolutionary game theory mostly account for additive effects of payoffs on fitness [33-36] or effects of payoffs on carrying capacity [43-45], overlooking the cross effect of payoffs and density on fitness; thereby the increased competition reduces or negates the selective advantage of altruism from assortment mechanisms [20-24]. Since density-dependent regulation and population fluctuation are fundamental characteristics of natural systems, the evolution of altruism through this route may be common in nature.

Since the average growth rate of altruists is only slightly higher than that of selfish individuals (Fig. 3A), the invasion of altruism necessarily experiences a long transient phase where altruists remain at a low density, followed by an abrupt regime shift to altruistic dominance (Fig. 1, Fig. S2 and S3). Unlike regime shifts caused by gradual changes in parameters near tipping points [46-48], this shift occurs without any parameter changes [49, 50]. The timing of the regime shift can be predicted analytically (inequality 3), marking the transition from a population dominated by selfish individuals to one dominated by altruists (Fig. 1, Fig. S2 and S3). Classical theory for variable environments typically assesses whether rare mutants can invade a resident population by comparing their long-term growth rates (the Lyapunov exponents), where invasion is expected if the mutant’s long-term growth rate exceeds that of the pre-invasion resident population [41, 42]. However, this approach is not applicable to our models, as the long-term growth rates of both initially rare altruists and selfish individuals are zero. Despite this, we find that altruists can invade when their short-term average growth rate over the past *t* generations surpasses that of selfish individuals, as indicated by 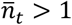. This short-term cumulative advantage can lead to the invasion success of altruism. Although the long-term average population density of a fluctuating population may remain unchanged (Fig. 2A), the short-term average can vary over time (Fig. 4A-F). For the regime shift to occur, these short-term fluctuations must exceed a threshold (inequality 2, Fig. 2-5). Rare altruistic mutants initially incur high costs that boost the density of selfish individuals, causing the average population density to surpass the threshold (inequality 2 and 5). Once this happens, altruists begin to outcompete and competitively exclude selfish individuals (Fig. 4D-I, Fig. 5). Evidently, the early increase in population density is almost entirely driven by selfish individuals, not altruists. While altruists lose battles against selfish individuals in terms of immediate payoffs, they ultimately prevail by enduring the long transient phase and achieving dominance in the long run.

The traditional view suggests that the evolution of altruism must require some forms of preferential interactions among altruists (i.e., assortment). For example, kin selection theory posits that individuals carrying an altruistic gene tend to help their close relatives that are likely carrying the same gene [10-12]. Group selection theory suggests that small groups of altruistic individuals can be favored by natural selection [13, 14]. Reciprocal altruism theory ensures assortative interactions through complex iterative strategies [15-17], social tags [51-54] or specific population structures [55-57]. Essentially, these theories all prescribe the condition of Hamilton’s rule to ensure that the average payoff of altruists exceeds that of selfish individuals, either in classical [10, 11] or nonlinear form [12, 58] or under variable environments [59, 60]. These altruistic behaviours described by Hamilton’s rule can be interpreted as instances of narcissistic altruism [19]. In contrast, we show that altruists can invade and dominate a selfish population under fluctuating selection triggered by increased competition from payoffs, without relying on assortment mechanisms, provided that population dynamics fluctuate above the threshold set by the invasion criterion (inequality 2 and 5). Such fluctuations can arise from intrinsic complex dynamics (Fig. 1, Fig. S2 and S3), environmental stochasticity (Supplementary Material S3, Fig. S5), or demographic stochasticity (Supplementary Material S4, Fig. S7). The invasion criterion obliviates the constraint of Hamilton’s rule in all its forms [12, 18, 58-60] and enables altruism to evolve with a consistently lower average payoff than selfishness, thereby allowing the emergence of true altruism. Altruists can invade and extirpate selfish individuals even when the cost of altruism exceeds its resulting benefit (Fig. 3C and D, Fig. S3), suggesting the possibility for extremely self-sacrificial altruism.

In conclusion, true altruistic behaviour can emerge and persist without the need for assortment under fluctuating selection if competition is intensified by social benefits from the invasion of altruism. Fluctuating selection from payoff-regulated density dependence presents an alternative paradigm for explaining the evolution of altruism, shifting away from assortment mechanisms but toward the ecological contexts that drive complex population dynamics. While social experiments rarely test the effects of fluctuating selection, environmental variability has been shown to strongly predict the biogeographic distribution of cooperative breeding in birds [61], offering indirect evidence that social behaviour can evolve under fluctuating selection. Instead of forming alliances through assortment mechanisms, sacrificial altruism creates a temporal mirage of greater social benefits, fueling conflicts among opponents, especially in a volatile society. This mirrors the divide-and-conquer strategy often used in warfare and politics, where short-term loses are strategically endured to ultimately defeat a powerful adversary.

## Supporting information

This PDF file includes Supplementary Materials S1 to S5 and Figures S1 to S10.

## Acknowledgments

This work was supported by Xishuangbanna Tropical Botanical Garden (RC2023056BN) and the National Research Foundation of South Africa (89967).

## Author Contributions

F.Z., A.H. and C.H. conceived the research. F.Z. performed the research and wrote the paper. All authors contributed substantially to revisions.

