## Supplementary material for "Fluctuating selection breaks Hamilton’s rule for the evolution of altruism": This PDF file includes Supplementary Materials S1 to S5 and Figures S1 to S10.

Supplementary Materials S1 to S5
Figures S1 to S10

**S1. Derivation of the replicator equation from Eq. (1)**

The replicator equation is a fundamental theoretical framework in evolutionary biology that describes the dynamics of strategy frequencies [1]. It is often used to investigate evolutionary games. We here derive the replicator equation from Eq. (1) in the main text, and show Eq. (1) is an extension of the replicator equation by further incorporating the density dependence of population growth. If we remove the density-dependent term $1 - n_t$, Eq. (1) becomes

$$\begin{aligned} x_{t+1} &= x_t \exp(r + \lambda \xi_t)\,, \\ y_{t+1} &= y_t \exp(r + \lambda \zeta_t)\,. \end{aligned} \tag{S1}$$

Thus, we have

$$\frac{x_{t+1}}{x_{t+1} + y_{t+1}} = \frac{x_t \exp(r + \lambda \xi_t)}{x_t \exp(r + \lambda \xi_t) + y_t \exp(r + \lambda \zeta_t)} = \frac{\frac{x_t}{x_t + y_t} \exp(\lambda \xi_t)}{\frac{x_t}{x_t + y_t} \exp(\lambda \xi_t) + \frac{y_t}{x_t + y_t} \exp(\lambda \zeta_t)}, \tag{S2}$$

Let $p_t = x_t/(x_t + y_t)$ be the frequency of altruists in the population, and we can rewrite Eq. (S2) as

$$p_{t+1} = \frac{p_t \exp(\lambda \xi_t)}{p_t \exp(\lambda \xi_t) + (1 - p_t) \exp(\lambda \zeta_t)} = \frac{p_t}{p_t + (1 - p_t) \exp[\lambda(\zeta_t - \xi_t)]}. \tag{S3}$$

Note that Eq. (S3) is the classical replicator equation with discrete generations. As the average payoff of altruists is always lower than that of selfish individuals (i.e., $\zeta_t - \xi_t > 0$), the frequency of altruists will converge to zero, confirming that selfishness is the Nash equilibrium.

Similarly, we can derive the dynamical equation for the frequency of altruists directly from Eq. (1) in the main text. Calculating $x_{t+1}/(x_{t+1} + y_{t+1})$, we have

$$\frac{x_{t+1}}{x_{t+1} + y_{t+1}} = \frac{x_t \exp[(r + \lambda \xi_t)(1 - n_t)]}{x_t \exp[(r + \lambda \xi_t)(1 - n_t)] + y_t \exp[(r + \lambda \zeta_t)(1 - n_t)]}. \tag{S4}$$

Thus, we obtain

$$p_{t+1} = \frac{p_t \exp[(r + \lambda \xi_t)(1 - n_t)]}{p_t \exp[(r + \lambda \xi_t)(1 - n_t)] + (1 - p_t) \exp[(r + \lambda \zeta_t)(1 - n_t)]}, \tag{S5}$$

and then

$$p_{t+1} = \frac{p_t}{p_t + (1 - p_t) \exp[\lambda(\zeta_t - \xi_t)(1 - n_t)]}. \tag{S6}$$

Evidently, the convergence of Eq. (S6) is no longer determined solely by the positivity of $\zeta_t - \xi_t$, but is also affected by the term of $1 - n_t$. Notably, while the replicator equation (S3), derived from the two-dimensional system (S1), is a closed one-dimensional system, Eq. (S6) is not closed because it includes the variable $n_t$. Therefore, we need to derive the dynamic equation for the population density $n_t$. From Eq. (1) in the main text, we can obtain

$$n_{t+1} = x_{t+1} + y_{t+1} = x_t \exp[(r + \lambda \xi_t)(1 - n_t)] + y_t \exp[(r + \lambda \zeta_t)(1 - n_t)]\,, \tag{S7}$$

and then

$$n_{t+1} = n_t\{p_t \exp[(r + \lambda \xi_t)(1 - n_t)] + (1 - p_t) \exp[(r + \lambda \zeta_t)(1 - n_t)]\}. \tag{S8}$$

Thus, we get a closed two-dimensional system (combining Eq. S6 and S8) coupling the dynamics of altruists' frequency ($p_t$) and population density ($n_t$), which is equivalent to Eq. (1).

The classical replicator equation simply assumes that the subpopulation of each strategy increases at a rate proportional to the received average payoff, i.e., the growth rate of each strategy is affected additively by its average payoff. This assumption leads to Eq. (S1) and, consequently, the replicator equation (Eq. S3). According to Eq. (S1), the density of altruistic strategy can increase indefinitely, even though its frequency converges to zero according to Eq. (S3) (see Fig. S1). This occurs in the absence of density-dependent regulation on population growth. In this study, following the basic assumption of the replicator equation, we consider the more realistic ecological context of density-dependent regulation, which surprisingly leads to fluctuating selection and completely reversed the well-known expectation about the evolution of altruism (see the main text).

**S2. Derivation of inequality (2) and (3)**

From Eq. (1) in the main text, the relative growth rates of altruists and selfish individuals at generation $t$ are given by

$$\begin{aligned} \log\left(\frac{x_{t+1}}{x_t}\right) &= (r + \lambda\xi_t)(1 - n_t), \\ \log\left(\frac{y_{t+1}}{y_t}\right) &= (r + \lambda\zeta_t)(1 - n_t). \end{aligned} \tag{S9}$$

Since the population densities across generations are bounded, the long-term growth rate (the Lyapunov exponent) of the pre-invasion selfish population is zero [2], and can be calculated as

$$\lim_{t\to\infty} \frac{1}{\mathrm{t}} \sum_{i=0}^{t-1} \log\left(\frac{y_{t+1}}{y_t}\right) = \lim_{t\to\infty} \frac{1}{\mathrm{t}} \sum_{i=0}^{t-1} r(1 - n_t) = r\left(1 - \lim_{t\to\infty} \frac{1}{\mathrm{t}} \sum_{i=0}^{t-1} n_t\right) = r(1 - \tilde{n}) = 0, \tag{S10}$$

where $\zeta_t = 0$ and $\tilde{n}$ is the long-term average population density of the pre-invasion selfish population. From Eq. (S10), we have $\tilde{n} = 1$. Similarly, we can calculate the long-term growth rate of initially rare altruists as follows

$$\lim_{t\to\infty} \frac{1}{\mathrm{t}} \sum_{i=0}^{t-1} \log\left(\frac{x_{t+1}}{x_t}\right) = \lim_{t\to\infty} \frac{1}{\mathrm{t}} \sum_{i=0}^{t-1} (r - \lambda c)(1 - n_t) = \frac{r - \lambda c}{r} \lim_{t\to\infty} \frac{1}{\mathrm{t}} \sum_{i=0}^{t-1} r(1 - n_t) = 0, \tag{S11}$$

where $\xi_t = -c$ due to the rarity of initial altruists. Consequently, we cannot evaluate invasiveness of altruists by comparing the long-term growth rate of initially rare altruists to that of resident selfish population as in the classical theory for variable environments [2]. Therefore, we adopt their short-term growth rates to assess whether initially rare altruists can invade a selfish population.

From Eq. (S9), the time-averaging relative growth rates over $t$ generations for altruists and selfish individuals are

$$\begin{aligned} \frac{1}{t}\log\left(\frac{x_t}{x_0}\right) &= \frac{1}{t}\log\left(\frac{x_t}{x_{t-1}}\frac{x_{t-1}}{x_{t-2}}\cdots\frac{x_1}{x_0}\right) = \frac{1}{\mathrm{t}}\sum_{i=0}^{t-1}(r + \lambda\xi_i)(1 - n_i), \\ \frac{1}{t}\log\left(\frac{y_t}{y_0}\right) &= \frac{1}{t}\log\left(\frac{y_t}{y_{t-1}}\frac{y_{t-1}}{y_{t-2}}\cdots\frac{y_1}{y_0}\right) = \frac{1}{\mathrm{t}}\sum_{i=0}^{t-1}(r + \lambda\zeta_i)(1 - n_i), \end{aligned} \tag{S12}$$

and the difference in the time-averaging relative growth rate between altruists and selfish individuals is

$$\frac{1}{t}\log\left(\frac{x_t}{x_0}\right)-\frac{1}{t}\log\left(\frac{y_t}{y_t}\right)=\frac{1}{t}\sum_{i=0}^{t-1}\lambda(\xi_i-\zeta_i)(1-n_i)=\lambda c(\bar{n}_t-1), \qquad (S13)$$

where $\xi_i - \zeta_i = -c$ (see the main text) and $\bar{n}_t$ is the average population density over the $t$ generations.

Furthermore, from Eq. (1) in the main text, we can derive the following equation for the density ratio of altruistic to selfish individuals:

$$\frac{x_{t+1}}{y_{t+1}}=\frac{x_t}{y_t}\exp[\lambda(\xi_t-\zeta_t)(1-n_t)]\,. \qquad (S14)$$

Note that the term in square brackets is the change rate of the density ratio at generation $t$, which is also the difference in relative growth rate between altruistic and selfish individuals (i.e., the difference between both equations in Eq. S9). Solving Eq. (S14) for $x_t/y_t$, we have

$$\frac{x_t}{y_t}=\frac{x_0}{y_0}\exp\left[\sum_{i=0}^{t-1}\lambda(\xi_i-\zeta_i)(1-n_i)\right]=\frac{x_0}{y_0}\exp\left[\frac{1}{t}\sum_{i=0}^{t-1}\lambda(\xi_i-\zeta_i)(1-n_i)\,t\right]. \qquad (S15)$$

Noting that $(1/t)\sum_{i=0}^{t-1}\lambda(\xi_t-\zeta_t)(1-n_t)=\lambda c(\bar{n}_t-1)$ being the difference in the time-averaging relative growth rate between altruists and selfish individuals (see Eq. S13), we have the following equation

$$\frac{x_t}{y_t}=\frac{x_0}{y_0}\exp[\lambda c(\bar{n}_t-1)t]\,. \qquad (S16)$$

Evidently, the density of altruists increases relative to the density of selfish individuals (i.e., altruists can invade the selfish population) when $\lambda c(\bar{n}_t-1)>0$, leading to inequality (2) in the main text, i.e., $\bar{n}_t>1$.

Letting $x_t/y_t>1$ signal that altruists have taken over the population after the transient phase, from Eq. (S16), we have

$$\frac{x_0}{y_0}\exp[\lambda c(\bar{n}_t-1)t]>1, \qquad (S17)$$

which is equivalent to inequality (3) in the main text, i.e.,

$$\lambda c(\bar{n}_t-1)t>\log\left(\frac{y_0}{x_0}\right). \qquad (S18)$$

**S3. Ricker model with environmental stochasticity**

We introduce environmental noise into Eq. (1) in two distinct ways, with each based on different assumptions. If the per-capita growth rate of the population follows a normal distribution, the environmental noise can be considered in Eq. (1) as follows,

$$\begin{aligned} x_{t+1}&=x_t\exp[(r+\lambda\xi_t)(1-n_t)+\omega_t]\,,\\ y_{t+1}&=y_t\exp[(r+\lambda\zeta_t)(1-n_t)+\omega_t]\,, \end{aligned} \qquad (S19)$$

where $\omega_t{\sim}N(0,\sigma^2)$ is the normally distributed noise (experienced by both strategies due to being in the same population), with $\sigma$ representing the noise intensity. In contrast, if we assume that the population density follows a normal distribution, as per Noble et al. [3], the environmental noise can be represented in Eq. (1) as follows:

$$\begin{aligned} x_{t+1} &= x_t \exp[(r + \lambda\xi_t)(1 - n_t)]\,(1 + \omega_t), \\ y_{t+1} &= y_t \exp[(r + \lambda\zeta_t)(1 - n_t)]\,(1 + \omega_t). \end{aligned} \tag{S20}$$

The model of Eq. (S19) demonstrates a regime shift of altruism invasion, even in case where the corresponding deterministic system does not undergo population fluctuations (i.e., $r < 2$) (Fig. S5, Fig. S9C and D). On the other hand, model (S20) does not exhibit a regime shift, regardless of whether the corresponding deterministic system fluctuates (see Fig. S6, Fig. S9E and F). This is because the invasion criterion for the regime shift in these two stochastic models is still the average population density over generations being greater than one ($\bar{n}_t > 1$) (see the following proofs). However, model (S20) fails to meet this invasion criterion of altruism (see Fig. S6C and F).

In what follows, we prove $\bar{n}_t > 1$ is still a necessary condition (the invasion criterion) for the regime shift of altruism invasion in the models (S19) and (S20). From Eq. (S19), the time-averaging relative growth rates over $t$ generations for altruists and selfish individuals are

$$\frac{1}{t}\log\left(\frac{x_t}{x_0}\right) = \frac{1}{t}\log\left(\frac{x_t}{x_{t-1}}\frac{x_{t-1}}{x_{t-2}}\cdots\frac{x_1}{x_0}\right) = \frac{1}{t}\sum_{i=0}^{t-1}(r + \lambda\xi_i)(1 - n_i) + \frac{1}{t}\sum_{i=0}^{t-1}\omega_i, \tag{S21}$$

$$\frac{1}{t}\log\left(\frac{y_t}{y_0}\right) = \frac{1}{t}\log\left(\frac{y_t}{y_{t-1}}\frac{y_{t-1}}{y_{t-2}}\cdots\frac{y_1}{y_0}\right) = \frac{1}{t}\sum_{i=0}^{t-1}(r + \lambda\zeta_i)(1 - n_i) + \frac{1}{t}\sum_{i=0}^{t-1}\omega_i, \tag{S22}$$

Thus, the difference in the time-averaging relative growth rate between altruistic and selfish individuals can be calculated as

$$\frac{1}{t}\log\left(\frac{x_t}{x_0}\right) - \frac{1}{t}\log\left(\frac{y_t}{y_0}\right) = \frac{1}{t}\sum_{i=0}^{t-1}\lambda(\xi_i - \zeta_i)(1 - n_i) = \lambda c(\bar{n}_t - 1). \tag{S23}$$

Therefore, altruists can invade the selfish population when $\lambda c(\bar{n}_t - 1) > 0$, leading to $\bar{n}_t > 1$.

For the stochastic model (S20), using the same method as the above, we can estimate the time-averaging relative growth rates over $t$ generations for altruists and selfish individuals:

$$\frac{1}{t}\log\left(\frac{x_t}{x_0}\right) = \frac{1}{t}\sum_{i=0}^{t-1}(r + \lambda\xi_i)(1 - n_i) + \frac{1}{t}\sum_{i=0}^{t-1}\log(1 + \omega_i)\,, \tag{S24}$$

$$\frac{1}{t}\log\left(\frac{y_t}{y_0}\right) = \frac{1}{t}\sum_{i=0}^{t-1}(r + \lambda\zeta_i)(1 - n_i) + \frac{1}{t}\sum_{i=0}^{t-1}\log(1 + \omega_i)\,, \tag{S25}$$

Thus, the difference in the averaging relative growth rate between altruistic and selfish individuals is the same as Eq. (S23), and then altruists can invade the selfish population when $\bar{n}_t > 1$. Therefore, for model (S19) and (S20), the invasion criterion of altruism is still inequality (2) as in the main text, i.e., $\bar{n}_t > 1$.

### S4. Ricker model with demographic stochasticity

To incorporate demographic stochasticity into Eq. (1), the population density is rescaled to population size by multiplying a constant $k$ to reflect the system size. Consequently, the population size of generation $t + 1$ (denoted by $N_{t+1}$) can be drawn from a Poisson distribution with the mean

$$\mathbb{E}(N_{t+1}) = kn_t\{p_t \exp[(r + \lambda\xi_t)(1 - n_t)] + (1 - p_t)\exp[(r + \lambda\zeta_t)(1 - n_t)]\}, \tag{S26}$$

i.e., the right-hand side of Eq. (S8) multiplied by $k$. The number of altruists (denoted by $N_{C,t+1}$) is a random number drawn from a binomial distribution with parameters $N_{t+1}$ and

$$\frac{p_t \exp[(r+\lambda\xi_t)(1-n_t)]}{p_t \exp[(r+\lambda\xi_t)(1-n_t)] + (1-p_t)\exp[(r+\lambda\zeta_t)(1-n_t)]}, \qquad (S27)$$

i.e., the right-hand side of Eq. (S5). Assuming any strategy may mutate into the other during the reproduction of the next generation, with a small mutation rate $\mu$. If using $N'_{C,t+1}$ to denote the number of altruists after adjustment for mutation, the population density and the frequency of altruists in generation $t+1$ can be calculated by $n_{t+1} = N_{t+1}/k$ and $p_{t+1} = N'_{C,t+1}/N_{t+1}$, respectively. Our simulations also show that altruists can invade a selfish population under demographic stochasticity, regardless of whether the corresponding deterministic system fluctuates (see Fig. S7).

**S5. A discrete logistic model for the evolution of altruism**

Since the growth factors of the Ricker model, $x_{t+1}/x_t$ and $y_{t+1}/y_t$ in Eq. (1), are exponential functions, the convexity of such nonlinear functions could potentially play a role in generating and triggering the regime shift of altruism invasion. To investigate the impact of the convexity in the growth factors, we consider the following discrete logistic model for the evolution of altruism:

$$\begin{aligned} x_{t+1} &= \mathrm{x_t} + x_t(r+\lambda\xi_t)(1-n_t), \\ y_{t+1} &= \mathrm{y_t} + y_t(r+\lambda\zeta_t)(1-n_t), \end{aligned} \qquad (S28)$$

where the symbols have the same meanings as in Eq. (1). The growth factors in model (S28) are linear functions of the population density. This model also demonstrates a similar regime shift of altruism invasion (see Fig. S10), suggesting that the convexity of the growth factors is not a necessary cause for generating the regime shift.

In addition, the invasion criterion of altruism is no longer the average population density over generations being larger than one ($\bar{n}_t > 1$). To derive the invasion condition of altruism for model (S28), we estimate the time-averaging relative growth rates over $t$ generations for altruists and selfish individuals as follows:

$$\frac{1}{t}\log\left(\frac{x_t}{x_0}\right) = \frac{1}{t}\sum_{i=0}^{t-1}\log[1+(r+\lambda\xi_i)(1-n_i)], \qquad (S29)$$

$$\frac{1}{t}\log\left(\frac{y_t}{y_0}\right) = \frac{1}{t}\sum_{i=0}^{t-1}\log[1+(r+\lambda\zeta_i)(1-n_i)], \qquad (S30)$$

For convenience, let $M_i = (r+\lambda\xi_i)(1-n_i)$ and $N_i = (r+\lambda\zeta_i)(1-n_i)$. Using the second-order Taylor series $\log(1+x) \approx x - \frac{1}{2}x^2$, Eq. (S29) and (S30) can be approximated as:

$$\frac{1}{t}\log\left(\frac{x_t}{x_0}\right) = \frac{1}{t}\sum_{i=0}^{t-1}\left(M_i - \frac{1}{2}M_i^2\right) = \frac{1}{t}\sum_{i=0}^{t-1}(r+\lambda\xi_i)(1-n_i) - \frac{1}{2t}\sum_{i=0}^{t-1}M_i^2, \qquad (S31)$$

$$\frac{1}{t}\log\left(\frac{y_t}{y_0}\right) = \frac{1}{t}\sum_{i=0}^{t-1}\left(N_i - \frac{1}{2}N_i^2\right) = \frac{1}{t}\sum_{i=0}^{t-1}(r+\lambda\zeta_i)(1-n_i) - \frac{1}{2t}\sum_{i=0}^{t-1}N_i^2. \qquad (S32)$$

Thus, the difference in the time-averaging relative growth rate between altruistic and selfish individuals can be calculated as

$$\frac{1}{t}\log\left(\frac{x_t}{x_0}\right)-\frac{1}{t}\log\left(\frac{y_t}{y_0}\right)=\lambda c(\bar{n}_t-1)-\frac{1}{2t}\sum_{i=0}^{t-1}(M_i^2-N_i^2). \qquad (S33)$$

Therefore, letting the right-hand side of Eq. (S33) be greater than zero, we get the invasion condition of altruism in a selfish population for model (S28) as follows

$$\bar{n}_t>1+\Omega_t, \qquad (S34)$$

where

$$\Omega_t=\frac{1}{2\lambda ct}\sum_{i=0}^{t-1}(M_i^2-N_i^2)=\frac{1}{2ct}\sum_{i=0}^{t-1}[2r+\lambda(\xi_i+\zeta_i)](\xi_i-\zeta_i)(1-n_i)^2<0. \qquad (S35)$$

Therefore, altruism invasion in the selfish population requires that the average population density over generations exceeds the threshold, $1+\Omega_t$, which is less than one (see Fig. S10).

The result above relies on approximations using the second-order Taylor series: $\log\,(1+M_i)\approx M_i-\frac{1}{2}M_i^2$ and $\log\,(1+N_i)\approx N_i-\frac{1}{2}N_i^2$. It implies that the absolute values of $M_i$ and $N_i$ must be sufficiently small for inequality (S32) to hold. More specifically, by letting $\log(1+M_i)=M_i-\frac{1}{2}M_i^2+R_1(M_i)$ and $\log(1+N_i)=N_i-\frac{1}{2}N_i^2+R_2(N_i)$, where $R_1(M_i)=\sum_{k=3}^{\infty}\frac{(-1)^{k+1}M_i^k}{k}$ and $R_2(N_i)=\sum_{k=3}^{\infty}\frac{(-1)^{k+1}N_i^k}{k}$ are the remainder terms, inequality (S32) can be modified as

$$\bar{n}_t>1+\Omega_t+\frac{1}{\lambda ct}\sum_{i=0}^{t-1}[R_2(N_i)-R_1(M_i)]. \qquad (S36)$$

Thus, the threshold for altruism invasion can be accurately estimated by appropriately truncating the higher-order terms of the remainders.

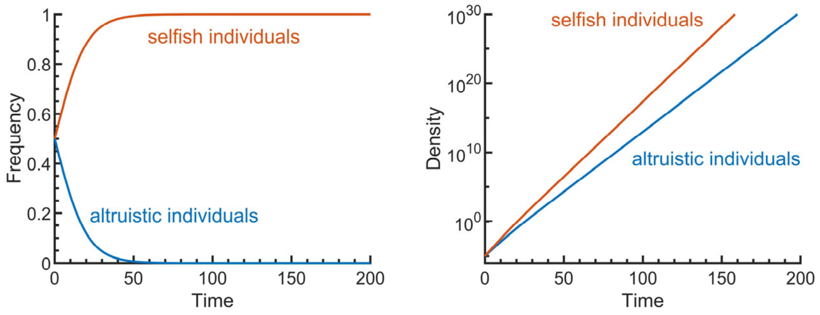


**Figure S1.** The frequency dynamics of the replicator equations for altruistic and selfish strategies (left panel; Eq. S3) and the density dynamics of the corresponding exponential growth equations (right panel; Eq. S1). Although the frequency of altruistic strategy converges to zero, its density still increases indefinitely. Parameter used are $b = 0.2$, $c = 0.1$, $\lambda = 1$, $r = 0.5$, $x_0 = 0.00001$, and $y_0 = 0.00001$.

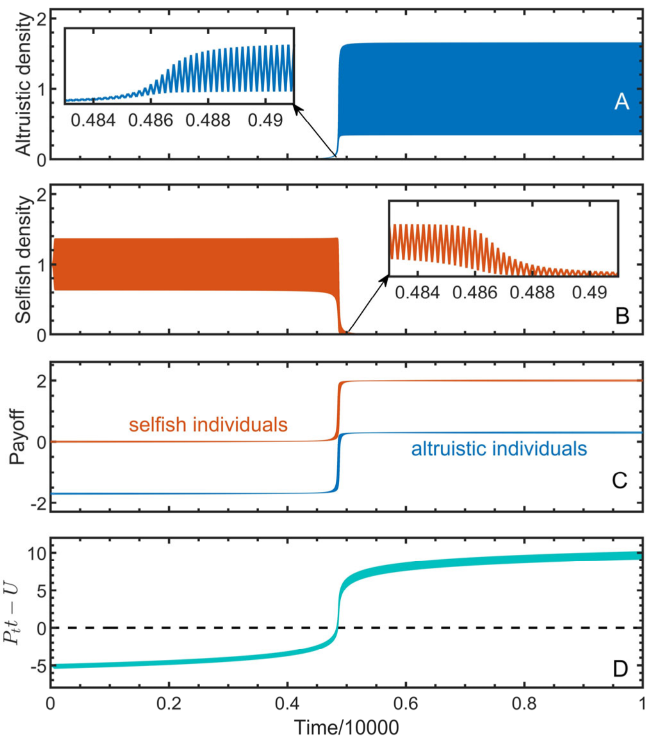


**Figure S2.** Emergence of altruism through a regime shift after a transient phase in a periodically fluctuating population. Panel (A) shows the dynamics of altruists starting from a small initial density in a selfish population, (B) shows the dynamics of selfish individuals, and (C) displays the average payoffs of altruistic and selfish individuals. Panel (D) demonstrates how the duration of the transient phase results from the inequality (3), where $P_t t$ and $U$ are the left- and right-hand side of the inequality, respectively. The parameters used are: $b = 2$, $c = 1.7$, $\lambda = 1$, $r = 2.1$, $x_0 = 0.001$, and $y_0 = 1$.

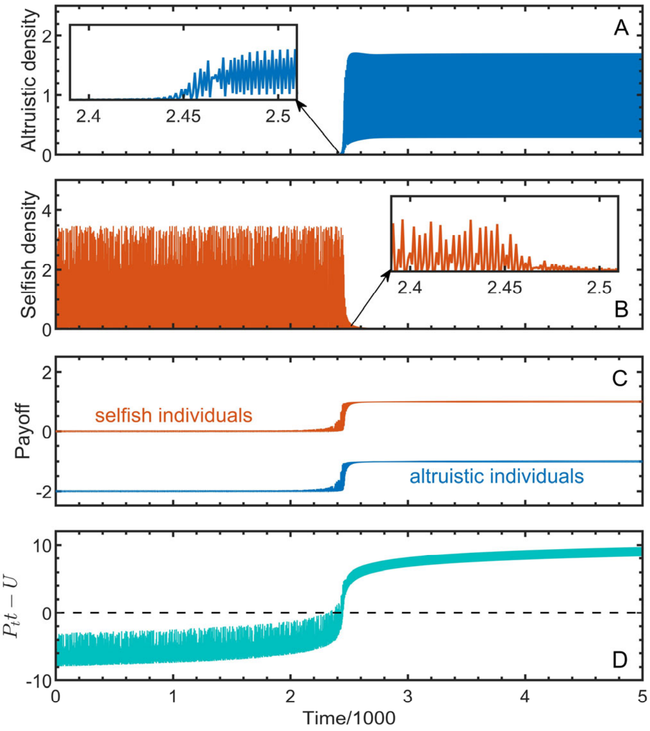


**Figure S3.** Emergence of altruism even when the cost is higher than the benefit. Panel (A) shows the dynamics of altruists starting from a small initial density in a selfish population, (B) shows the dynamics of selfish individuals, (C) displays the average payoffs of altruistic and selfish individuals. Panel (D) demonstrates how the duration of the transient phase results from the inequality (3), where $P_t t$ and $U$ are the left- and right-hand side of the inequality, respectively. The parameters used are: $b = 1$, $c = 2$, $\lambda = 1$, $r = 3.5$, $x_0 = 0.0001$, and $y_0 = 1$.

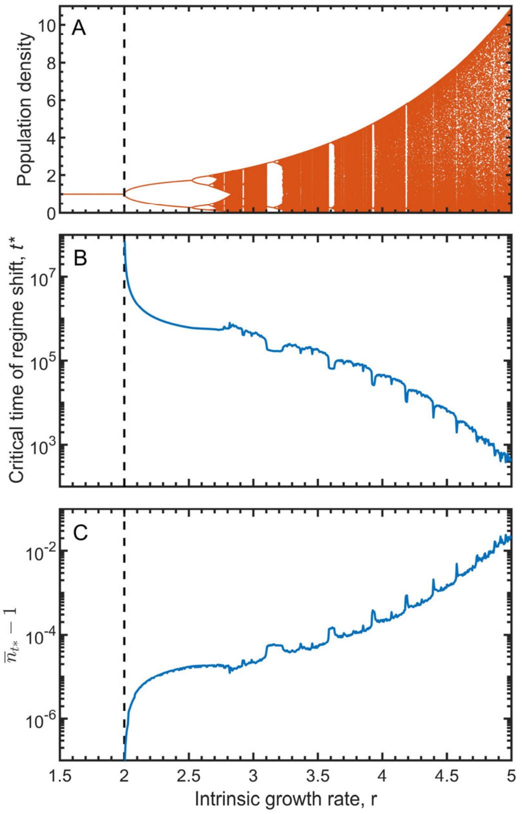


**Figure S4.** The regime shifts of altruism invasion in the case when $b < c$. Panel (A) shows the bifurcation diagram of the selfish population dynamics ($y_{t+1} = y_t \exp[r(1 - y_t)]$); (B) shows the dependence of the critical time of regime shift on the intrinsic growth rate; and (C) displays $\bar{n}_{t^*} - 1$, corresponding to the curves in panel (B), where $\bar{n}_{t^*}$ is the average population density over generations up to the critical time $t^*$. The parameters are: $b = 1$ and $c = 1.15$, $x_0 = 0.00001$ and $y_0 = 1$.

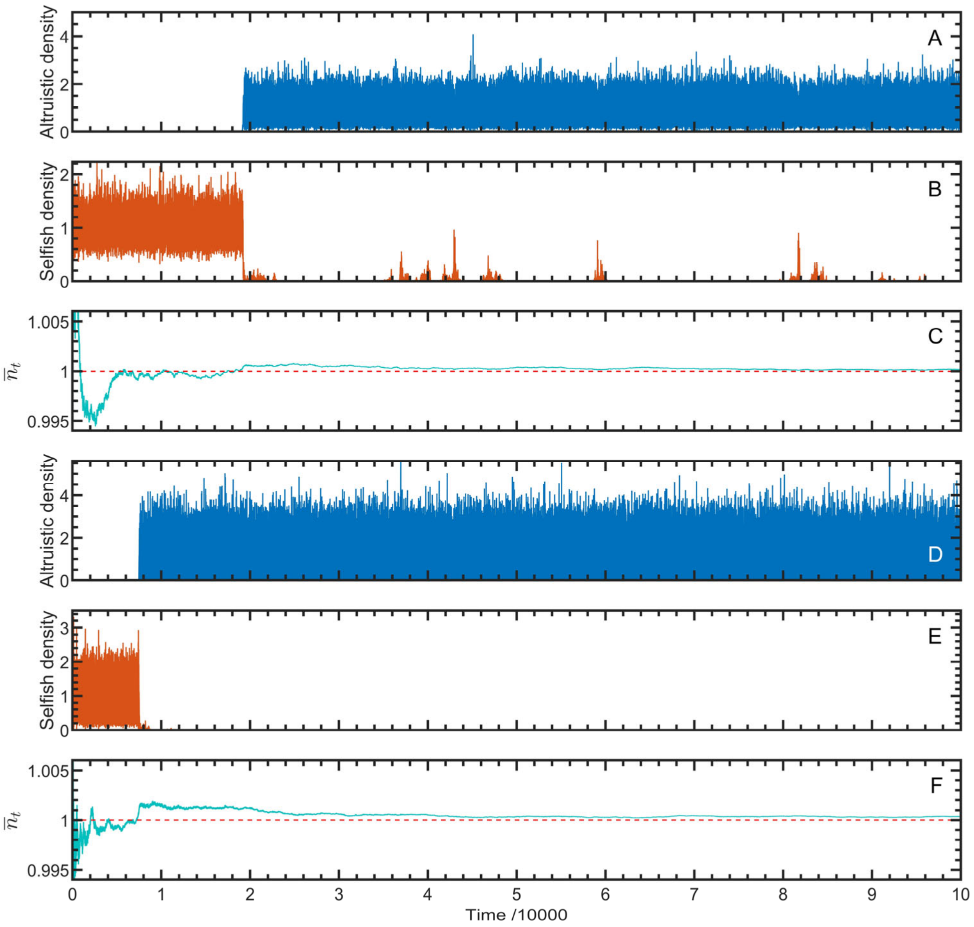


**Figure S5.** Emergence of altruism in noisy environments, as depicted by Eq. (S19). The regime shift of altruism invasion occurs regardless of whether the corresponding deterministic system fluctuates (from D to F) or not (from A to C). Parameters: $b = 2$, $c = 1.2$, $\lambda = 1$, $r = 1.5$ for (A to C) and $r = 2.3$ for (D to F); $x_0 = 0.0001$, $y_0 = 1$; $\sigma = 0.2$.

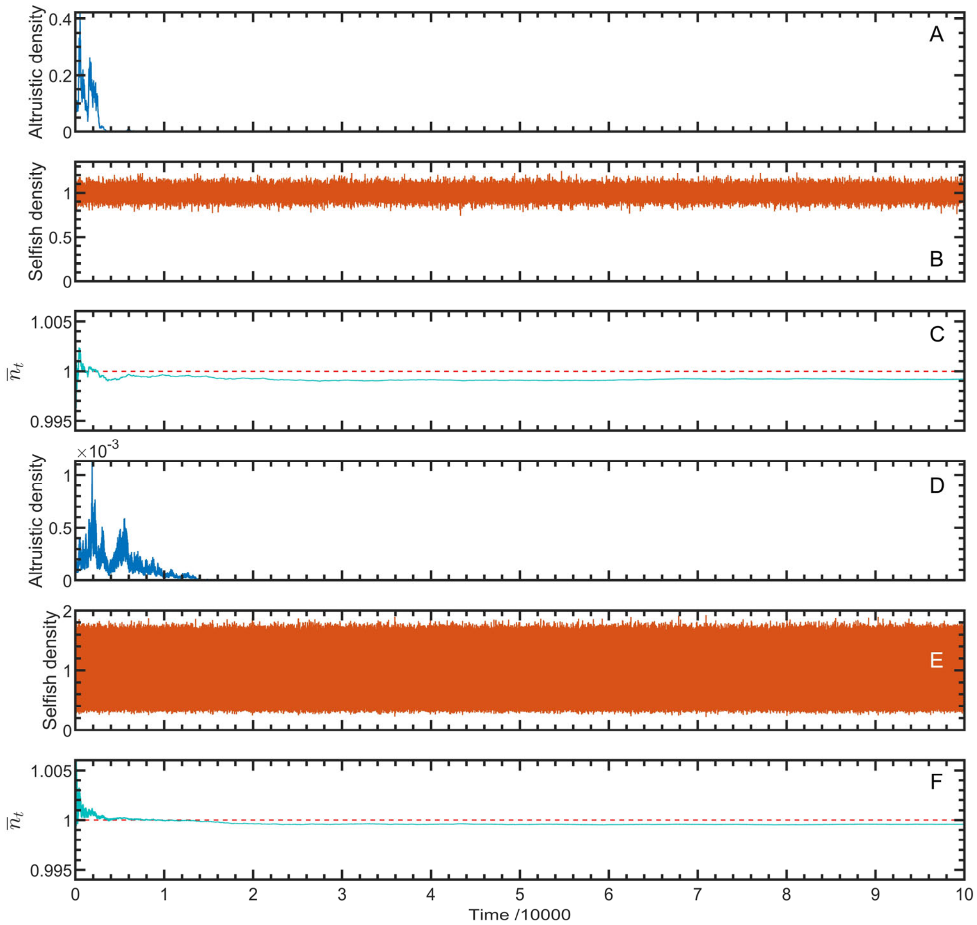


**Figure S6.** Altruism cannot invade selfish populations under the environmental noise described in Eq. (S20), regardless of whether the corresponding deterministic system fluctuates (from D to F) or not (from A to C). Parameters: $b = 2$, $c = 1.2$, $\lambda = 1$, $r = 1.5$ for (A to C) and $r = 2.3$ for (D to F); $x_0 = 0.0001$, $y_0 = 1$; $\sigma = 0.05$.

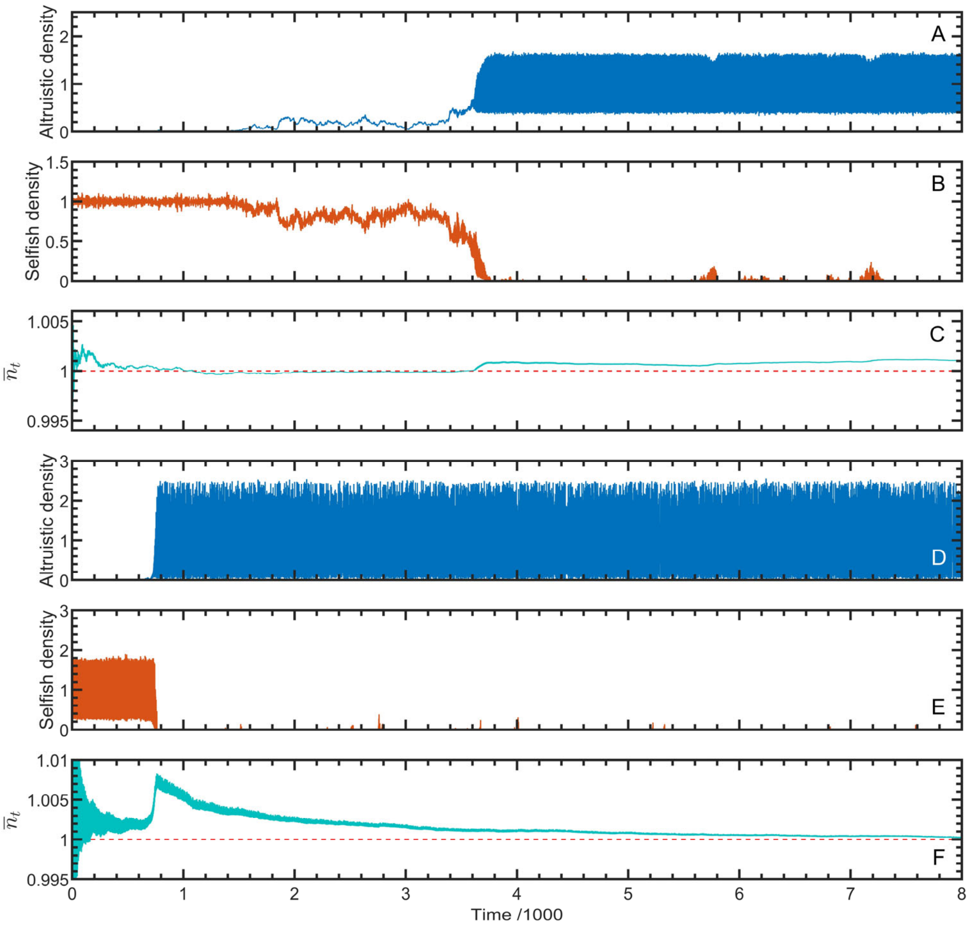


**Figure S7.** Emergence of altruism under demographic stochasticity, as depicted in Supplementary Material S4. The regime shift of altruism invasion occurs regardless of whether the corresponding deterministic system fluctuates (from D to F) or not (from A to C). Parameters: $b = 2$, $c = 1.5$, $\lambda = 1$, $r = 1.8$ for (A to C) and $r = 2.5$ for (D to F); $x_0 = 0$, $y_0 = 1$, $k = 2000$, $\mu = 0.00005$.

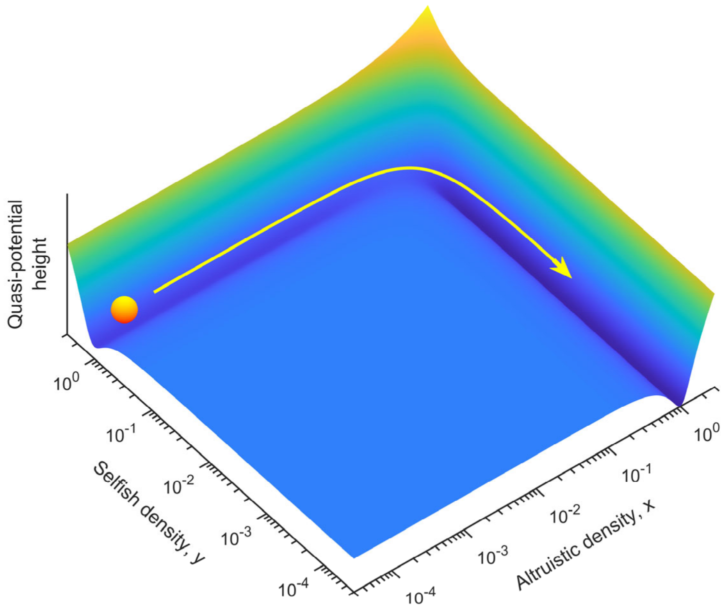


**Figure S8.** Illustration of a quasi-potential surface on the phase plane ($x$-$y$ space). The orange ball represents the evolving population, and the yellow arrow indicates the potential evolutionary direction of altruism invasion. A dynamical system can be visualized as a ball rolling on a quasi-potential surface over the phase plane, with wells corresponding to locally stable attractors and peaks unstable ones. Depending on its initial position, the ball will roll towards the bottom of a local well with the rate of convergence proportional to the slope of the quasi-potential surface. If the slope around a well is relatively flat, the rolling ball will move rather slowly, representing a long transient phase leading towards the local well. For the population described by Eq. (1), the points on the line $x + y = 1$ $(0 \leq x \leq 1)$ are all equilibria, forming a ditch on the quasi-potential surface. The deepest point corresponds to (1,0), representing a pure altruistic population, followed by (0,1), representing a pure selfish population, and the other points on the line have relatively high potential but are still at the bottom of the ditch, representing a population with mixed strategies. Although the system can stand still at any point (equilibrium) on this line, its position is prone to disturbance. Population fluctuations can induce the ball to shift position successively along the ditch, leading to the regime shift of altruism invasion (comparing Fig. S9A with B). Disturbances such as environmental noise can also drive the ball away from a well, prolonging transient phenomena and even triggering a regime shift (see Fig. S9C and D). These transient dynamics differ from stable asymptotic behaviours but can flicker for a long period before shifting to alternative attractors [4].

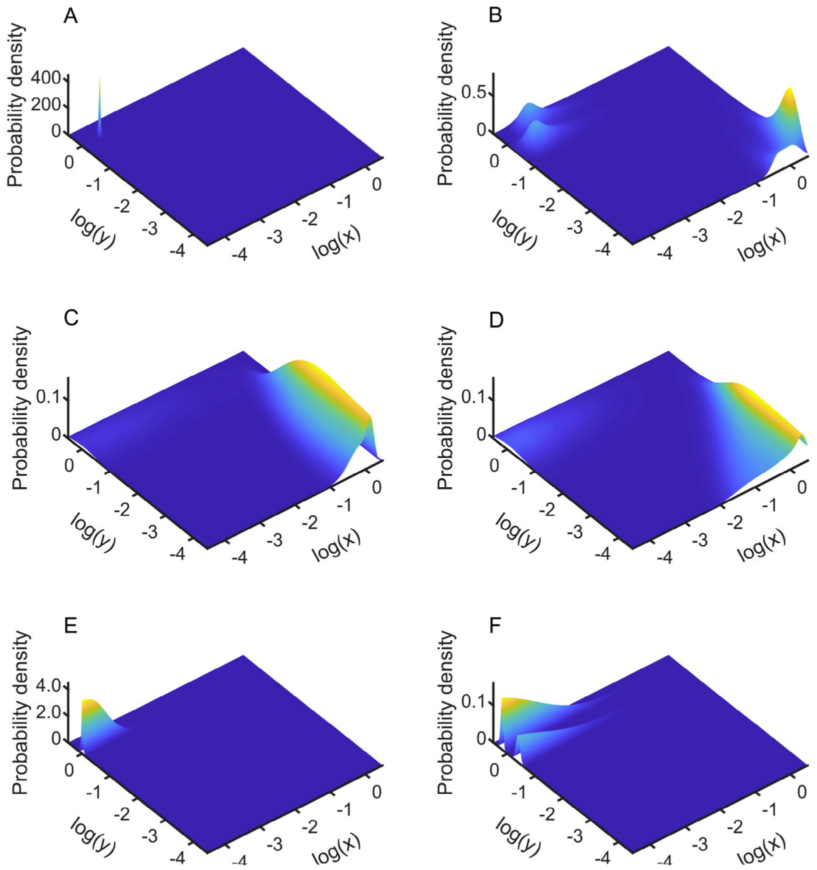


**Figure S9.** Joint probability distribution of strategy densities. The surface is estimated using a 2-dimensional kernel density method from multiple simulations of altruism invasion in a selfish population, conceptually illustrated on the quasi-potential surface in Fig. S8. The regime shifts from selfishness to altruism occurred in B to D but not in A, E and F. Panel A and B shows the probability density estimated from simulations of 200,000 generations of Eq. (1) starting from $x_0 = 0.0001$ and $y_0 = 1$, and panel (C to F) estimated from 100 repeated stochastic simulations starting from $x_0 = 0.0001$ and $y_0 = 1$, with each running 100,000 generations. C and D used the stochastic models: $x_{t+1} = x_t \exp[(r + \lambda\xi_t)(1 - n_t) + \omega_t]$ and $y_{t+1} = y_t \exp[(r + \lambda\zeta_t)(1 - n_t) + \omega_t]$, where $\omega_t \sim N(0, \sigma^2)$ being noise (the same for both strategies) with $\sigma = 0.2$ representing the noise intensity; E and F ran the models $x_{t+1} = x_t \exp[(r + \lambda\xi_t)(1 - n_t)](1 + \omega_t)$ and $y_{t+1} = y_t \exp[(r + \lambda\zeta_t)(1 - n_t)](1 + \omega_t)$ but with $\sigma = 0.05$ (see Supplementary Material S3, Fig. S5 and S6). Parameters: $b = 2$, $c = 1.2$, $\lambda = 1$, $r = 1.5$ for (A, C, E) and $r = 2.3$ for (B, D, F).

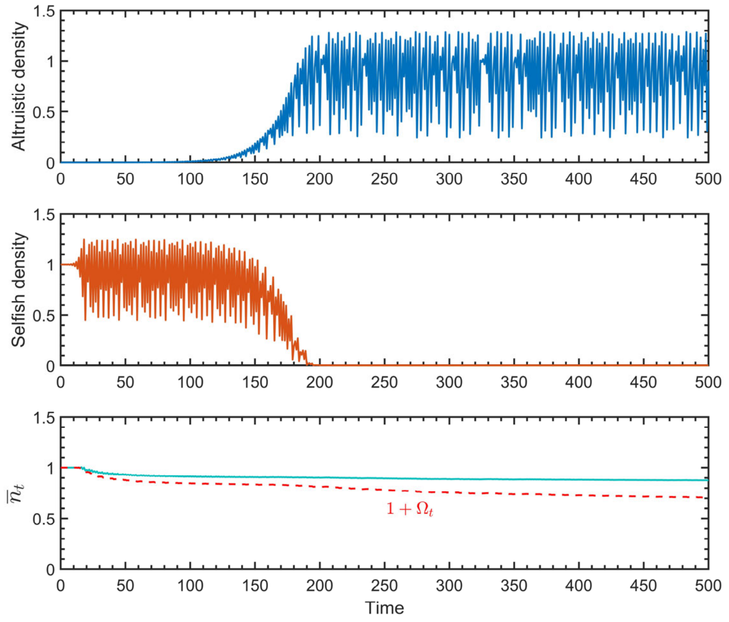


**Figure S10.** The regime shift of altruism invasion in the discrete logistic model, as described by Eq. (S28), with linear growth factors. The cyan curve in the bottom panel represents the average population density, which exceeds the invasion threshold, $1 + \Omega_t$, indicated by the red dashed line (see Supplementary Material S5). Parameters: $b = 1$, $c = 0.8$, $\lambda = 1$, $r = 2.6$, $x_0 = 0.0001$, $y_0 = 1$.
